# Exposure to titanium dioxide nanoparticles accelerates abnormal fat deposition through the mediation of the ROS via the AGE-RAGE signaling pathway in mice

**DOI:** 10.1101/2023.08.24.554746

**Authors:** Jingyi Liu, Liang Zhong, Tiantian Yuan, Jingchun Sun, Yaxin Wang, Jiahao Chen, Gongshe Yang, Taiyong Yu

**Affiliations:** College of Animal Science and Technology, Northwest A&F University, No.22 Xinong Road, Yangling, Shaanxi 712100. China

**Keywords:** Titanium dioxide nanoparticles, Nano-safety, Fat deposition, Adipocyte, AGE-RAGE, ROS

## Abstract

Titanium dioxide nanoparticles (TiO_2_ NPs) are widely added to various types of foods as food additives. Previous studies have shown that TiO_2_ NPs exposure can cause abnormal deposition of adipose tissue in organisms, resulting in lipid metabolism disorder. However, the potential molecular mechanisms underlying TiO_2_ NPs’ effects have yet to be elucidated. In this study, our data indicated that TiO_2_ NPs (100 mg/Kg BW, 20 nm) accelerated abnormal fat deposition in the epididymal adipose tissues and disturbed the level of blood glucose subsequently in normal-fat diet mice. Further studies showed that TiO_2_ NPs at a concentration of 100 µg/mL significantly induced the proliferation and differentiation of 3T3-L1 preadipocytes. Mechanistic studies we revealed that TiO_2_ NPs induced the overproduction of reactive oxygen species (ROS) in the cytoplasm, which subsequently upregulated the expression of receptors for advanced glycation end products (AGEs) leading to lipid accumulation in the adipocytes with a higher level of ROS. However, the antioxidant N-acetylcysteine (NAC) was a therapeutic potential for lipid overaccumulation in the adipocytes. This study provides insight into the mechanism underlying fat deposition induced by TiO_2_ NPs and highlighted the need for reevaluation of food-grade TiO_2_ NPs exposure in daily life.

## 1. Introduction

The applications of titanium dioxide (TiO_2_) permeate every aspect of our daily life. Owing to its brightness and high refractive index, TiO_2_ is widely used in food, medicine, paint, and other projects (Vance et al., 2015). In the past decades, food-grade TiO_2_ (commonly known as E171) is regarded as an inert material with good biosafety (EFSA Panel on Food Additives and Flavourings (FAF), et al., 2019). In daily life, TiO_2_ is often added to chocolate to increase its texture (Grande & Tucci, 2016;), or used in foods with high lipid content, such as cream cake, candy, and carbonated drinks, and for making all types of foods more attractive (Weir et al., 2012; He et al., 2022).

TiO_2_ is composed of micron-sized particles (MP_S_) and nano-sized particles (NP_S_). Compared with MPs, nano-sized TiO_2_ has a larger specific surface area and interacts with cells, thereby affecting cell secretion and metabolic processes and functions (Shi, Magaye, Castranova, & Zhao, 2013). Although the gastrointestinal absorption rate of TiO_2_ NPs is low and 99.9%-99.98% of orally administered TiO_2_ NPs are excreted in the feces, they can accumulate in living organisms (EFSA Panel on Food Additives and Flavourings (FAF), et al., 2021; Fu et al., 2020). It has been shown that these TiO_2_ NPs with diameters less than 100 nm can be deposited in the liver, kidney, brain, spleen, reproductive organs and other organs via blood circulation, causing adverse effects on the organs and systems of living organisms (Shakeel et al., 2016). The safety of TiO_2_ NPs has been questioned, and there has been widespread concern about the environmental and health problems caused by them (Luo et al., 2020). Currently, TiO_2_ is no longer considered a safe food additive, and on January 14, 2022, the European Commission banned the use of TiO_2_ (E171) as a food additive (EFSA Panel on Food Additives and Flavourings, 2021; Luo et al., 2020).

With the improvement in the living standards of people, glucose and lipid metabolism disorders, such as obesity, diabetes, and cardiovascular diseases, are becoming increasingly common. Excessive intake of high sugar and high-fat diet (HFD) will induce obesity, and it is worth noting that these foods often contain TiO_2_ NPs. Some early reports confirming the toxicity of TiO_2_ NPs have shown that exposure to TiO_2_ NPs can increase blood glucose in mice by inducing endoplasmic reticulum stress in the liver, and has significant insulin tolerance (Han et al., 2020; Hu et al., 2016; Hu et al., 2018). On this basis, treatment of mice with both antioxidants (resveratrol and vitamin E) significantly reduced the increase in reactive oxygen species (ROS) due to TiO_2_ NPs and inhibited the rise in blood glucose (Hu et al. 2016). Another study showed that oral administration of TiO_2_ NPs (30 ± 7 nm) significantly increased body weight, liver weight, and the amount of adipose tissue, especially in HFD-fed mice, by destroying the mucus layer and altering the intestinal microbiota (Zhu et al., 2021). It was found that pre-exposure to TiO2 NPs (200mg/kg) would aggravate alcohol related liver injury, liver steatosis and inflammation, and promote the development of Alcoholic liver disease (Liu et al., 2021). It has also been shown that exposure to TiO_2_ NPs (50 mg/kg BW) for 30 or 90 days can lead to significant liver edema, steatosis and necrosis (Chen et al., 2015). At the same time, Chen et al. showed by gavage of Sprague-Dawley rats with TiO_2_ NPs (29±9 nm) that significant changes in lipid histological characteristics in the serum of rats treated with TiO_2_ NPs (50 mg/kg BW). The accumulation of lipid peroxidation products (malondialdehyde, MDA) and a decrease in the activity of the antioxidant enzyme SOD were also observed and were closely associated with differential metabolites (Chen et al., 2020; Chen et al., 2022). Although the intake of TiO_2_ NPs has been shown to lead to disturbed glucose metabolism and lipid deposition in organisms, the underlying molecular mechanisms of TiO_2_ NPs-induced lipid deposition largely remain to be elucidated.

In this study, we investigated the biological effects of oral intake of TiO_2_ NPs (20 nm) on normal-fat diet mice by determining the fat deposition in various adipose tissues and other metabolic parameters *in vivo*, as well as their *in vitro* effects on 3T3-L1 preadipocytes and murine primary adipocytes. Mechanistically, proteomic sequencing analysis was applied to probe the related pathways under the exposure of TiO_2_ NPs to the adipocytes, and thus to further assess to reveal the importance of food-grade TiO_2_ NPs exposure, especially in the normal population.

## 2. Materials and methods

### 2.1 Characterization of TiO_2_ nanoparticles

Nano-TiO_2_ (99.9%, anatase, 20 nm, CAS:13463-67-7) was purchased from Deke Daojin Science and Technology Co., Ltd (Beijing, China). The primary particle sizes and morphologies were measured using scanning electron microscopy (SEM, Nano SEM-450, Czech) and transmission electron microscopy (TEM, HT7800, Japan), respectively, and the crystalline shape was determined using X-ray diffraction (XRD, Bruker D8 ADVANCE, Germany). After ultrasonic wave stirring for 30 min, the hydrodynamic diameter and zeta potential of TiO_2_ NPs in phosphate-buffered saline (PBS) were measured using a combined dynamic light scattering/particle electrophoresis instrument (Malvern Zetasizer Nano ZS series, UK).

### 2.2 Animals and treatments

Seven-week-old C57BL/6 male mice were purchased from the Xi’an Yifengda Biotechnology Co. Ltd and were acclimated to the environment for one week. Next, 10 mice of similar weight were selected randomly for each group. One group was exposed to 100 mg/kg BW of TiO_2_ NPs solution through oral gavage (TiO_2_ NPs powder was accurately weighed and dissolved in sanitary saline solution in an ultrasonicated bath for 30 min before gavage) while the mice in the other group were administered 10 mL/kg BW sanitary saline solution. Both groups of mice were treated for 12 weeks and fed a standard rodent chow diet (14049 Cooperative Medical Biological Engineering Co. Ltd., China). The mice were housed under a 12-h light/dark cycle and given *ad libitum* access to food and water during the experiment. All animal protocols were approved by the Committee of Experimental Animal Management at Northwest A & F University, China (2011-31101684).

Oral glucose tolerance test (OGTT) and insulin tolerance test (ITT) were performed on mice at weeks 9 and 11 postgavage, respectively. For the OGTT, the mice were fasted for 16 h and they maintained normal water intake during the fast before the oral administration of glucose (2.0 g/kg BW). Plasma glucose concentrations were measured at 15, 30, 60, 90, and 120 min for each mouse from collected tail vein blood. For the ITT, the mice were fasted for 4 h and they maintained normal water intake during the fast before the intraperitoneal injection of insulin solution (0.25 U/kg BW). All plasma glucose concentrations were determined using blood glucose test strips (Yuwell Medical Equipment Co. Ltd, China), and after the experiments, all the mice were supplemented with mice chow for recovery.

### 2.3 Blood and tissue sample collection

After the exposure to TiO_2_ NPs for 12 weeks, the mice were sacrificed using ether. Blood samples were collected and the serum was separated through centrifugation at 3000 rpm/min for 10 min at 4 °C. The serum samples were sent to Wuhan Servicebio Co. Ltd. for the detection of blood biochemical indicators, namely high-density lipoprotein (HDL), low-density lipoprotein (LDL), triacylglycerol (TG), total cholesterol (CHO), alanine aminotransferase (ALT), and aspartate aminotransferase (AST). Then inguinal white adipose tissues (iWAT), epididymal white adipose tissues (eWAT), brown adipose tissues (BAT), and various organs, such as the liver, lung, and spleen, were collected and weighed. All the samples were snap-frozen in liquid nitrogen and stored at −80 °C for further research.

### 2.4 Cell culture

3T3-L1 cells (Stem Cell Bank, Chinese Academy of Science) were cultured in Dulbecco’s modified Eagle’s medium (DMEM; Servicebio, China) supplemented with 10% fetal bovine serum (FBD; ZETA Life, China). For cell proliferation, the cells were treated with different concentrations of TiO_2_ NPs (0, 25, 50, and 100 µg/mL), and when the cell density reached approximately 30% after two cell cycles (36 h), they were collected for further experiments (Xi et al., 2019). For cell differentiation, 2 days postconfluence, the 3T3-L1 cells (designated day 0) were incubated in a cocktail medium (10% FBS, 0.5 mM 3-isobutyl-1-methylxanthine, 1 μM dexamethasone, and 5 μg/mL insulin) for 2 days (Xu et al., 2021). Next, the 3T3-L1 cells were transferred into DMEM containing 10% FBS and 5 μg/mL insulin for another 6 days of treatment. During the induction of 3T3-L1 differentiation, different concentrations of TiO_2_ NPs were added to the medium, and then the cell samples were collected on day 8 for further experiments.

Murine primary adipocytes were isolated from the iWAT of 3-week-old mice as previously reported (Chen et al., 2021), and then the cells were cultured in a growth medium (GM; DMEM/F12 with 10% FBS). Induction to differentiation was performed 2 days postconfluence of the primary adipocytes as described previously. Next, TiO_2_ NPs at a concentration of 100 µg/mL and N-acetyl-L-cysteine (NAC, CAS: 616-91-1, Aladdin) at a concentration of 10 mM were added to the medium for exposure to the cells as the design of experiments. All the cells were cultured at 37 °C in a humidified atmosphere with 5% CO_2_.

### 2.5 Detection of cell proliferation

Briefly, 10 μL of CCK-8 reagent (Solarbio, China) was added to each cell culture well and incubated for 2 h at 37 °C in a 5% CO_2_ incubator. After incubation, the absorbance of each well was measured at 650 nm as the reference wavelength and 450 nm as the detection wavelength (PerkinElmer, USA).

The cells were seeded and after 36 h of exposure, the cells were collected and washed twice with cold PBS (4 °C) and then fixed in 70% cold ethanol overnight at 4 °C. Subsequently, the fixed cells were stained with 50 mg/mL of DAPI for 30 min above the ice. Finally, the number of cells at different periods was determined using a flow cytometer (Becton Dickinson, USA).

Cells were cultured in a 96-well plate and transfected when the cell density reaches about 30%. The EdU staining was performed as described in the manual of the EdU cell proliferation detection kit (RiboBio, China).

### 2.6 Detection of cell differentiation

The 8-day differentiated cells were stained with the Oil Red O staining solution after washing off the residual medium with PBS. After staining, the images of lipid droplets were observed under a Nikon TE2000 microscope. In addition, isopropanol was applied to extract the Oil Red O stain, and quantitative analysis was performed at a wavelength of 510 nm.

The 8-day differentiated cells were washed with PBS to remove the medium, fixed in 4% polymethanol for half an hour, and stained with Bodipy dye for 30 min. The images of the lipid droplets were observed under a fluorescence microscope (Nikon, Japan).

### 2.7 Protein extraction and TMT labeling

Proteomics sequencing services were provided by Lianchuan Biotechnology Co. Ltd. (Zhejiang, China). The 3T3-L1 preadipocytes were treated with NC or 100 µg/mL TiO_2_ NPs 2 days postconfluence. Then, after another 6 days of differentiation, the cells were collected. Protein sample was added an appropriate amount of lysate (8m urea, 1% protease inhibitor Cocktail) and sonicated on the ice to promote its digestion. The sample was centrifuged at 20000g at 4 ℃ for 10min, and the supernatant was collected. The protein concentration was determined with BCA kit. Furthermore, used a TMT kit to mark protein according to the manufacturer’s protocol.

### 2.8 HPLC fractionation and LC−MS/MS analysis

The tryptic peptides were fractionated into fractions by high pH, a Waters XBridge Shield C18 RP column (3.5μm particles, 4.6 mm ID, 250 mm length) was used for HPLC fractionation. Each pre-separated fraction was dissolved in liquid phase A, centrifuged at 20000 g for 2 min, and the supernatant was transferred. Each fraction was analyzed by LC−MS/MS analysis for 1 h. The resulting MS/MS data were processed using the Maxquant search engine (v.1.5.2.8).

### 2.9 Measurement of reactive oxygen species (ROS) level in the cytoplasm

Intracellular ROS were determined using a ROS assay kit (Solarbio Life Science, CA1410, China). Briefly, the cells for further staining were washed three times with a serum-free medium. Meanwhile the ROS fluorescent probe DCFH-DA was diluted at 1:1000 with serum-free medium to a final concentration of 10 µmol/L and then added to the culture wells for 20 min in a 37 °C incubator. The cells were washed again three times with a serum-free medium to remove the DCFH-DA fully. Finally, the fluorescence intensity was determined using the Synergy H1 Hybrid MultiMode Reader (BioTek, Winooski, USA).

### 2.10 Real-time quantitative PCR (RT- qPCR)

Total RNA was extracted from the tissues or cells using TRIzol reagent (AG RNAex Pro Reagent, China). An amount of 1 μg RNA was reverse transcribed into cDNA using the PrimeScript RT Reagent Kit (Takara Biomedical Technology, China). For RT-qPCR, SYBR Green (Vazyme, China) was used with the StepOne Plus system (ABI, MA, USA). The 2^−ΔΔCt^ method was used to estimate the relative expression levels. All primers used for RT- qPCR are listed in Table S1.

### 2.11 Western blot analysis

Briefly, 10 μg of protein was extracted from the tissues or cells using the radioimmunoprecipitation assay lysis buffer (Beyotime, China). Total cellular proteins were separated using 10% SDS-PAGE gel and immunoblotted with the primary antibodies to PPARγ (ImmunoWay, YT3836), HSL (ImmunoWay, YT2240), LPL (Santa Cruz, sc-32382), ATGL (Cell Signaling, #2138), FABP4 (Abways, CY6768), FASN (Abcam, ab22759), CEBPα (ImmunoWay, YT0551), CyclinD1 (Abways, CY5404), PCNA (Cell Signaling, #2586) CDK4 (Abways, AB3183), p21 (Abways, CY5088), Nox4 (ABclone, A11274), RAGE (Abways, CY6633), p53 (Abways, CY5131), β-Actin (CWBIO, #50503), and β-tubulin (Sungene, KM9003).

### 2.12 Statistical analysis

All results from the cell- and mice-based experiments were at least three biological replicates. The data were initially collected in Excel and then imported into GraphPad Prism 7.0 software (GraphPad Software, San Diego, USA) for analysis and statistical plotting. All data are presented as the mean ± SEM. Two groups were compared using the Student’s *t*-test while multiple groups were compared using one-way ANOVA with Tukey’s post hoc test. The assumption of normality was tested using the Shapiro–Wilks test. All statistical tests were two-tailed, and statistical significance was considered at * *P* < 0.05, ** *P* < 0.01, *** *P* < 0.001, ns means not significant.

## 3. Results

### 3.1 Physicochemical properties of TiO_2_ nanoparticles

The physicochemical properties of inorganic nanoparticles, such as crystal structure, shape, size, and agglomeration, have a great effect on their biological actions. The TiO_2_ NPs used in this study were characterized in Fig 1 and S1. The XRD pattern in the whole spectrum of 2 theta values ranging from 30-50 showed that the TiO_2_ NPs used in this study were mainly in anatase form (Fig 1A). The TEM results indicated that the TiO_2_ NPs were mostly 20 nm in size (Fig 1B). In addition, the TEM and SEM images showed that the TiO_2_ NPs were prone to be in higher agglomeration state (Figs S1A and B). The hydrodynamic diameter and zeta potential for TiO_2_ NPs in PBS solution were 295.31 nm (Fig S1C) and -0.5626 mV (Fig S1D), respectively. However, the TEM images of the 3T3-L1 preadipocytes, which were treated with TiO_2_ NPs (diluted in PBS solution and added to a cell culture medium) showed that TiO_2_ NPs were absorbed and localized in the cytoplasm (Fig 1C).

**Fig 1.**
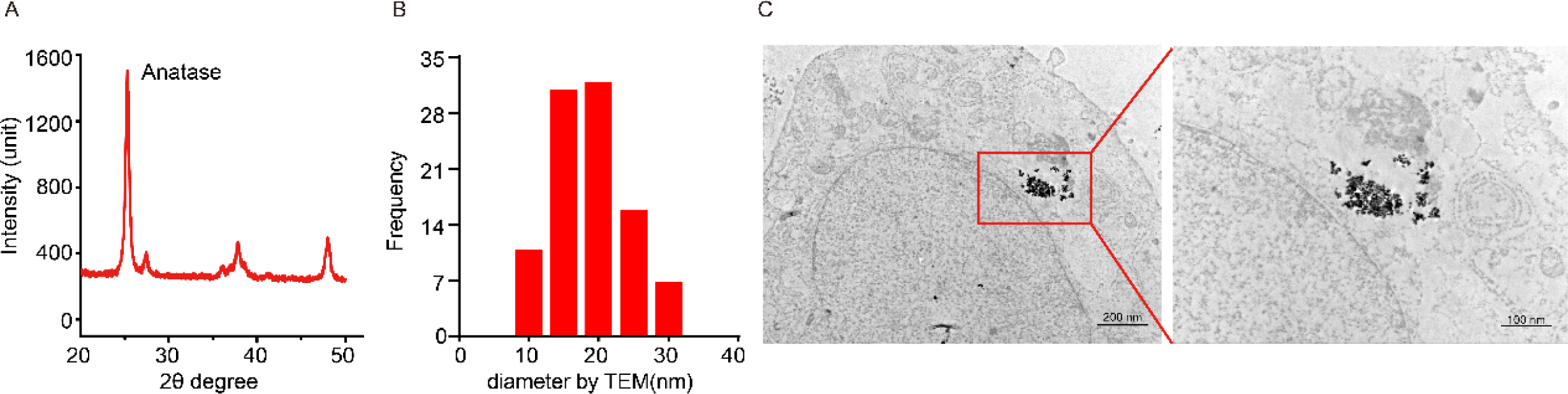
Physicochemical properties of titanium dioxide particles used in this study. (A) Crystal structure of TiO_2_ NPs. (B) The size distribution of TiO_2_ NPs by TEM. (C) TEM images of 3T3-L1 cells under the exposure of TiO_2_ NPs at different magnifications (Left 4000×, Right 8000×).

### 3.2 TiO_2_ NPs administration leads to increased blood glucose and fat accumulation in the eWAT in mice

A subchronic toxicity study design was applied to determine the biological effects of TiO_2_ NPs on normal-fat diet nonobese mice. We divided 20 C57BL/6J mice into two groups (Vehicle: n = 10; TiO_2_ NPs: n = 10) and fed them with a normal-fat diet (Fig 2A). After 12 weeks of TiO_2_ NPs treatment, the mice in both groups showed no significant effect on body weight gain or food intake (Fig 2B and C). However, the results of the OGTT showed that treatment with TiO_2_ NPs increased blood glucose significantly compared with that in the vehicle group (*P* < 0.001) while the area under the curve is significantly higher (Fig 2D, *P* < 0.05) for the TiO_2_ NPs group than for the vehicle group. The insulin ITT also showed that administration of TiO_2_ NPs resulted in reduced insulin sensitivity (Fig 2E). Subsequently, all the mice were sacrificed to collect the samples for analysis. The results indicated that the weight of the different adipose tissues is heavier in the TiO_2_ NPs-treated group, especially the eWAT expanded significantly (*P* < 0.01) than those in the vehicle group (Figs 2F and G). However, there was no significant difference in organ indexes, namely the heart, liver, spleen, lungs, and kidneys between the two groups (Fig 2H).

**Fig 2.**
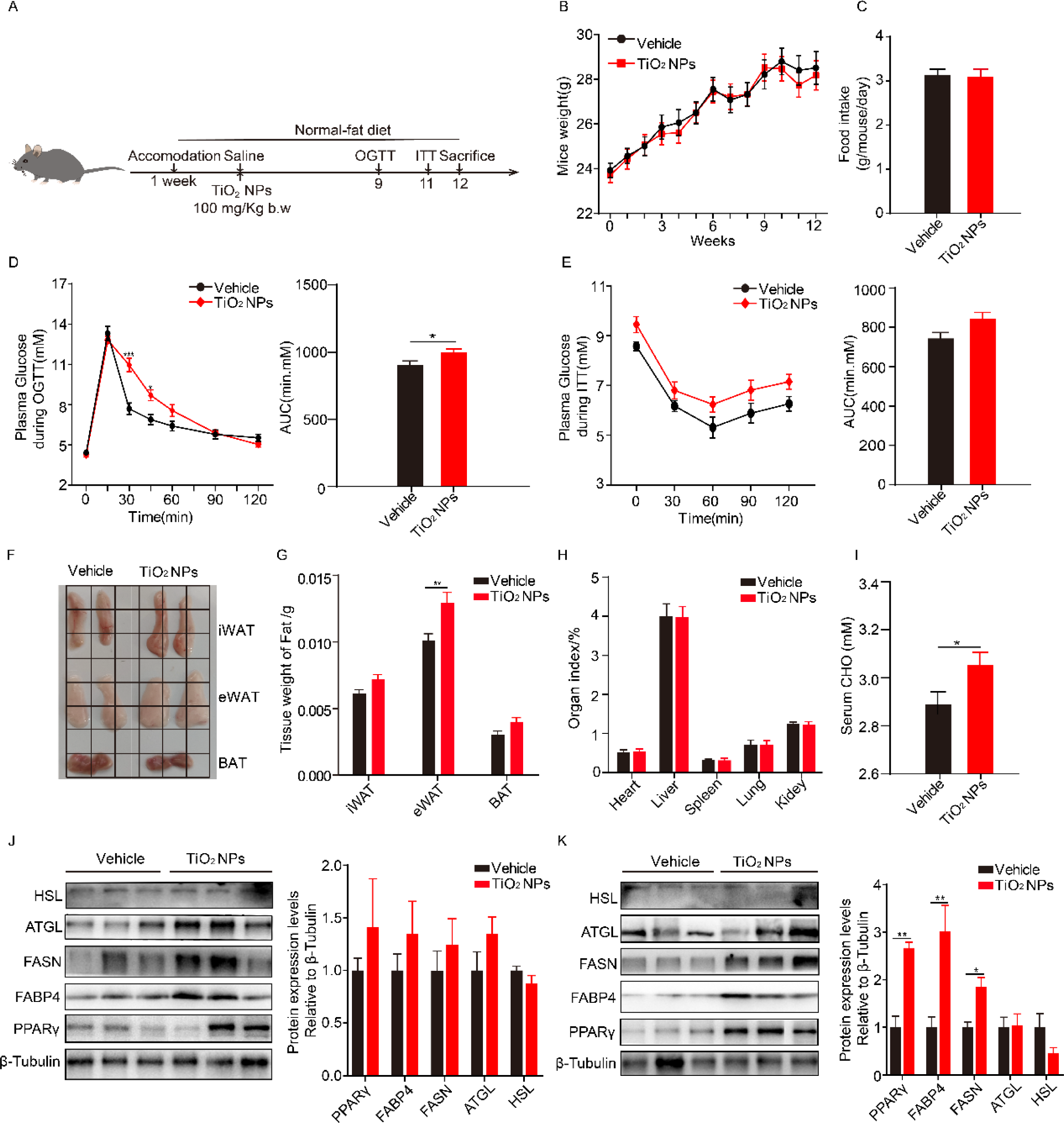
TiO_2_ NPs administration led to increased blood glucose and fat accumulation of epididymal adipose tissue in mice. (A) Schematic diagram of the experiment in mice. (B) The growth curve of body weight in mice (n = 10). (C) Statistical results of feed intaking in mice (n = 10). (D) OGTT in mice treated with TiO_2_ NPs or vehicle (n = 10). (E) ITT in mice treated with TiO_2_ NPs or vehicle (n = 10). (F) Representative images of different adipose tissues in mice. (G) Statistics fat weight of mice (n = 10). (H) Diagram of the organ index of mice (n = 10). (I) Serum total cholesterol (CHO) level (n = 10). (J) Western blot of the relative lipogenesis and lipolysis genes in iWATs (n = 3). (K) Western blot of the relative lipogenesis and lipolysis genes in eWATs (n = 3). Data represent mean ± standard error, \**P*<0.05, ** *P*< 0.01 and \*\*\**P* < 0.001 compared to the vehicle group.

### 3.3 TiO_2_ NPs administration shows no liver damage but alters the serum lipid levels in mice

To evaluate the toxicity of TiO_2_ NPs on mice livers, we measured the changes in several serum biochemical parameters. As shown in Fig S2, the levels of ALT and AST were not significantly different between the two groups (Fig S2A and B). Furthermore, the results of several biochemical parameters on serum lipid metabolism indicate that there are no significant changes (Figs S2C, D and E), excluding total cholesterol, which showed significantly higher levels (*P* < 0.05) in the TiO_2_ NPs group than that in the vehicle group (Fig 2I).

### 3.4 TiO_2_ NPs alter the expression of genes and protein levels in the various adipose tissues in mice

Based on the results, we determined the relative expression of genes, which correlates with lipid metabolism in the various adipose tissues. As presented in Fig S3, compared with the vehicle group, the relative mRNA levels of *FASN* and *HSL* in the iWAT in the TiO_2_ NPs-treated group increased significantly (*P* < 0.01) while the expression level of *SREBP1c* decreased significantly (Fig S3A, *P* < 0.01). Despite that, the expression levels of related genes in the eWAT are distinct from those of genes in the iWAT, and the expression levels of only *FASN* and *SREBP1c* increased significantly (*P* < 0.01) while the other related genes show no significant changes (Fig S3B). Furthermore, the results of the related genes in the BAT showed no significant difference between the two groups (Fig S3C). To determine the protein levels of related genes, we conducted western blotting on various WAT. The results showed no significant changes in the relative protein levels of PPARγ, FABP4, and FASN in the iWAT (Fig 2J) while the levels of the lipogenesis-related proteins including PPARγ, FABP4, and FASN are significantly (*P* < 0.01) elevated in the eWAT (Fig 2K).

### 3.5 TiO_2_ NPs induce preadipocyte proliferation and promote differentiation

To further verify the results from the mice treated with TiO_2_ NPs *in vivo* and to establish a cell model for mechanistic study *in vitro*, we chose the murine 3T3-L1 preadipocytes as the model for follow-up studies. Different doses of TiO_2_ NPs diluted in PBS solution were added to the cell culture medium. As shown in Fig 3A, though the CCK8 test showed no significant change in cell viability under each dose applied to the adipocytes, the EdU detection revealed an extraordinary variation where the number of EdU-positive cells increased significantly under a higher dose of TiO_2_ NPs (Fig 3B). In addition, the flow cytometry results also indicated that treatment of the preadipocytes with 100 µg/mL TiO_2_ NPs significantly (*P* < 0.05) decreased the number of cells in the G2/M (Figs 3C and D). Moreover, TiO_2_ NPs exposure to the preadipocytes led to a significant (*P* < 0.01) increase in the mRNA and protein levels of CylinD1, CDK4, and PCNA, and a significant decrease in the level of p21 (Figs 3E and F).

**Fig 3.**
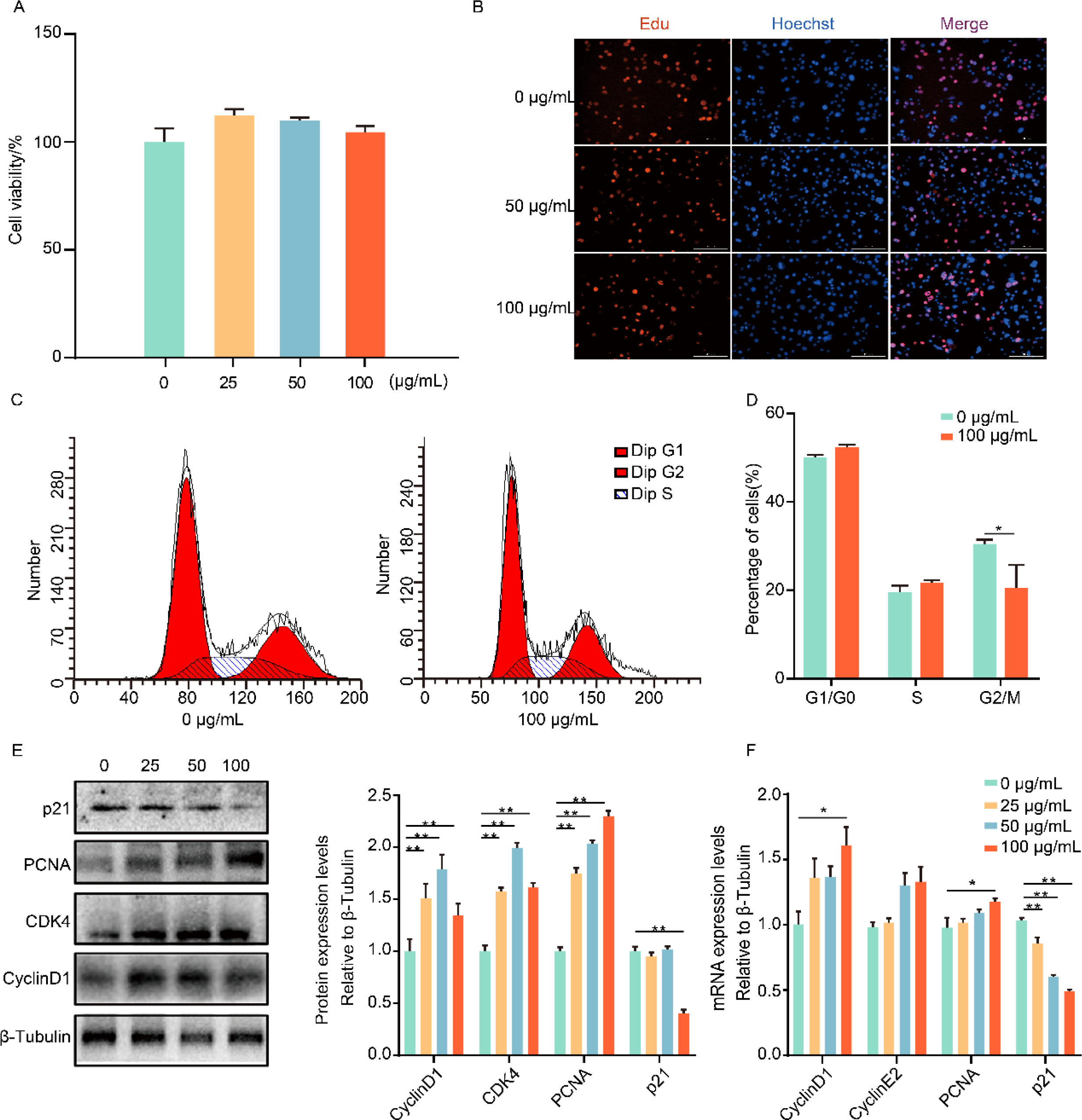
Effects of different doses of TiO_2_ NPs on the proliferation of 3T3-L1 preadipocytes. (A) CCK-8 test. (B) Images of EdU staining. (C) Flow cytometry results of preadipocytes. (D) Statistical results of flow cytometry. (E) Western blot of the relative marker genes of proliferation under the treatment of different doses of TiO_2_ NPs. (F) mRNA expression levels of the relative marker genes of proliferation. Data represent mean ± standard error, (n = 3), \**P*<0.05, ** *P*< 0.01 and \*\*\**P* < 0.001 compared to 0 µg/mL group.

To explore the roles of TiO_2_ NPs in the 3T3-L1 preadipocytes for cell differentiation, we conducted experiments using the same doses as those applied in the experiments on proliferation. From Fig 4A, we observed that the lipid accumulation detected using the Oil Red O was significantly (*P* < 0.01) potentiated under the management of higher doses of TiO_2_ NPs (50 and 100 µg/mL). In addition, the detection of Bodipy showed similar results (Fig 4B). Next, we found that the protein levels of the lipogenic genes (PPARγ and FABP4) increased significantly (*P* < 0.01), and the levels of the lipolytic genes (HSL) decreased significantly under the higher two doses of TiO_2_ NPs (50 and 100 µg/mL) (Fig 4C). The mRNA and protein levels of the related genes showed that the mRNA level of *CEBPα* significantly increased and the level of *HSL* significantly (*P* < 0.01) decreased (Fig 4D).

**Fig 4.**
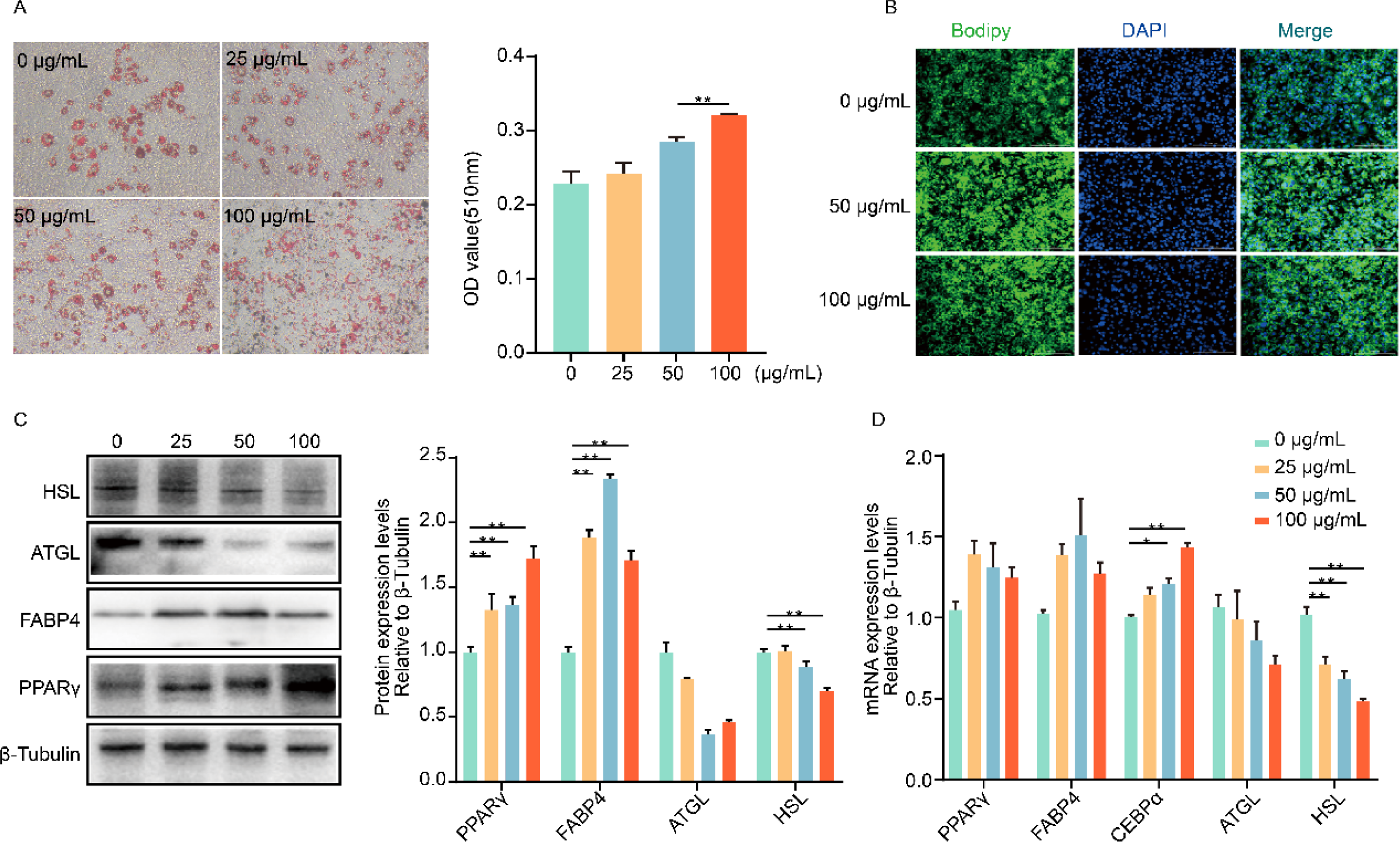
Effects of different doses of TiO_2_ NPs on the differentiation of 3T3-L1 preadipocytes. (A) Images of Oil Red O staining and quantitative statistics, magnification ×100. (B) Images of Bodipy staining. (C) Western blot of the relative marker genes of differentiation under the treatment of different doses of TiO_2_ NPs. (D) mRNA expression levels of the relative marker genes of differentiation. Data represent mean ± standard error, (n = 3), \**P*<0.05, ** *P*< 0.01 and \*\*\**P* < 0.001 compared to 0 µg/mL group.

### 3.6 Identification of related pathways associated with the differentiation of the 3T3-L1 preadipocytes using proteomic analysis

The specific mechanisms of the *in vitro* model are still unknown. Following the results obtained from the preadipocytes, we conducted quantitative research using liquid chromatography-tandem mass spectrometry analysis to uncover the mechanisms of TiO_2_ NPs that facilitate lipid deposition in the adipocytes (Fig 5A). Principal component analysis (PCA) showed good quantitative repeatability for the tested samples (Fig 5B). Then, approximately 5875 proteins were identified after mass spectrometry analysis, among which 4964 were quantifiable (Fig 5C). Next, the quantifiable proteins were filtered with the threshold fold change of > 1.2 and significance at *P* < 0.05, which yielded 1182 differentially expressed proteins, and 550 were upregulated and 632 were downregulated (Fig 5D).

**Fig 5.**
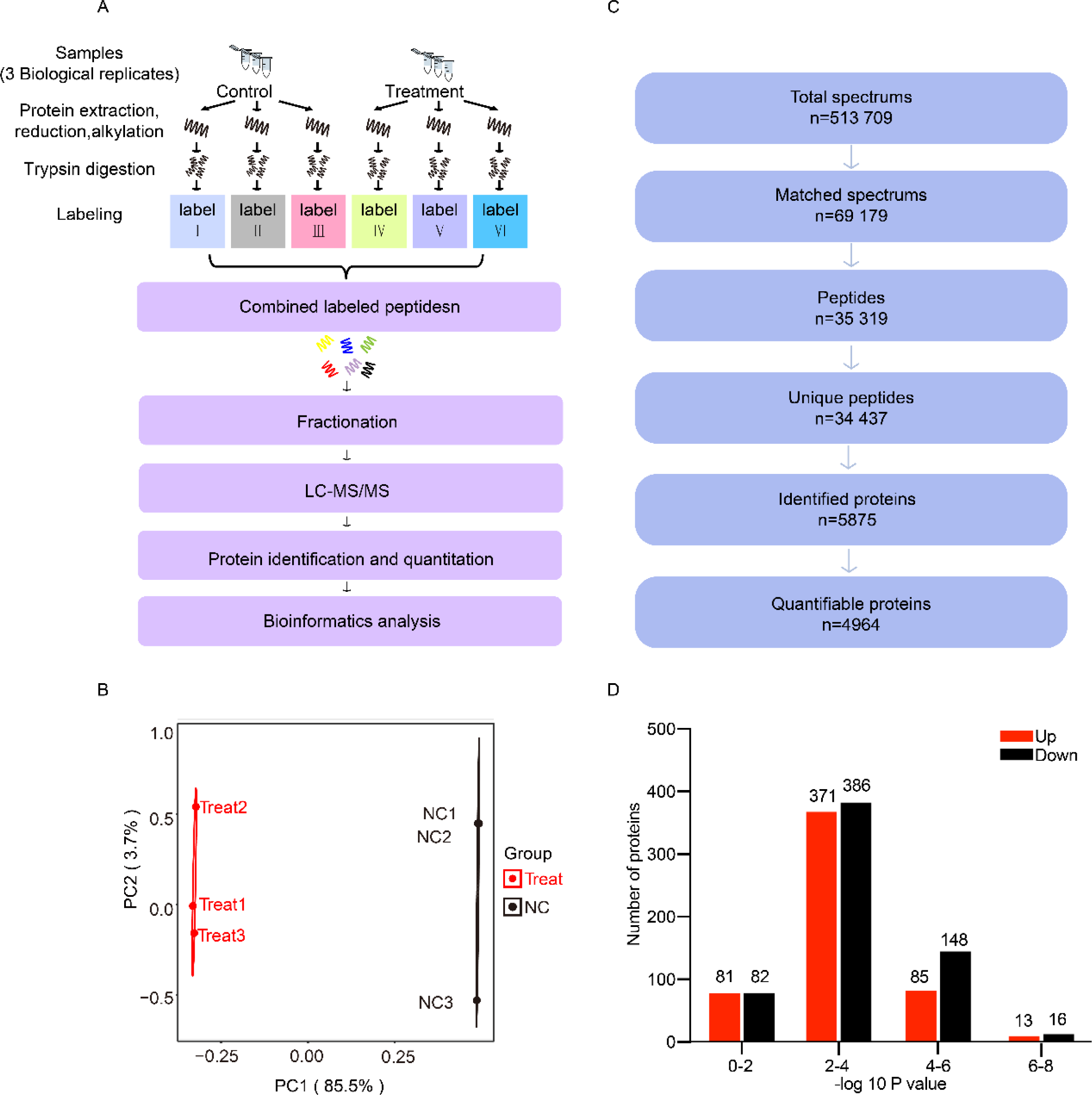
Identification of differentially expressed proteins using Proteomic Analysis. (A) Flow chart of the technical route for quantitative analysis of the treated group and control group. (B) Principal component analysis diagram of all sequenced samples. (C) LC−MS/MS spectral database search and analysis summary, including total spectra and effective spectra, the number of peptides, the number of unique peptides, the number of identifiable proteins, and the number of quantifiable proteins. (D) Statistical diagram of the number of differentially expressed proteins. Data are represented as mean± SEM (n = 3). Significance was determined by t-test analysis, \**P* < 0.05, \*\**P* < 0.01, and \*\*\**P* < 0.001.

As shown in the results in Fig 6A, most of the differentially expressed proteins were located in the cytoplasm (776, 49.9%) where the TiO_2_ NPs majorly functioned. Furthermore, the results of the Gene Ontology analysis revealed that there were other interesting biological processes, including the response to biotic stimulus and regulation of immune response except for the regulation of the lipid metabolic process (Figs S5A, B and C). Then, the molecular function displayed several optional ideas, such as signaling receptor activator activity, cytokine activity, and extracellular matrix structural constituent. The Kyoto Encyclopedia of Genes and Genomes (KEGG) pathway enrichment analysis showed that various typical pathways containing the p53 signaling pathway, as well as the AGE-RAGE signaling pathway, participated in the induction of adipocyte differentiation under the treatment of TiO_2_ NPs (Fig 6B).

**Fig 6.**
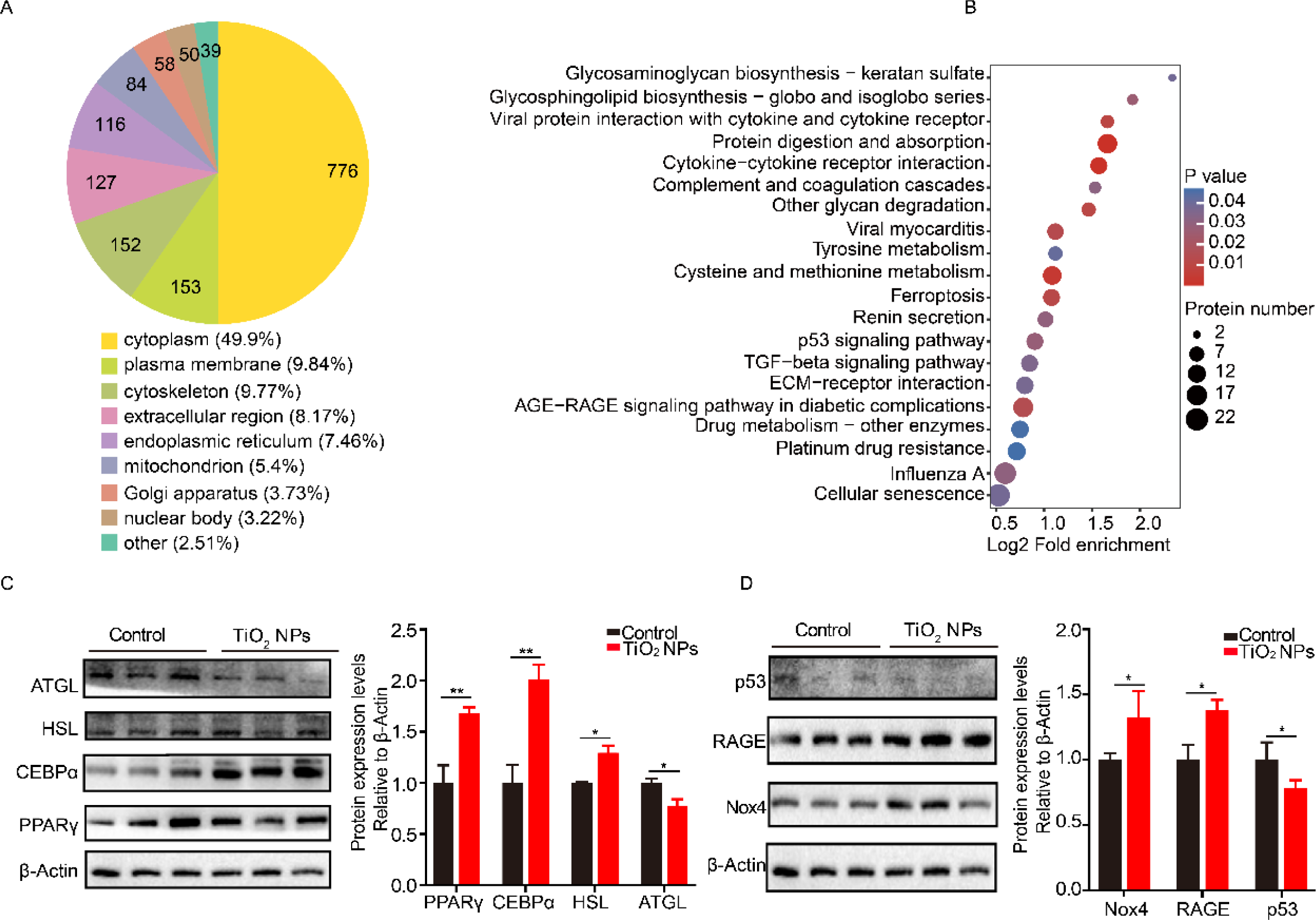
Functional analysis of differentially expressed proteins and effects of TiO_2_ NPs on the expression levels of relative genes in the mice primary adipocytes. (A) Subcellular localization of differentially expressed proteins. (B) KEGG pathway enrichment of differentially expressed protein. (C) Western blot of the related genes of lipogenesis and lipolysis in primary adipocytes. (D) Western blot of the related pathway genes in primary adipocytes. Data represent mean ± standard error, (n = 3), \**P*<0.05 ** *P*< 0.01, \*\*\**P* < 0.001 compared to the control group.

### 3.7 TiO_2_ NPs exacerbate lipid accumulation and upregulate the expression levels of related pathway genes in murine primary adipocytes

To reveal the potential mechanisms of TiO_2_ NPs function during adipogenesis, we isolated the primary white adipocytes from 3-week-old mice for further experiments. As the results indicated in the 3T3-L1 preadipocytes, the relative protein levels and the western blotting images showed that treatment of the primary white adipocytes with TiO_2_ NPs (100 µg/mL) significantly increased (*P* < 0.01) the protein levels of PPARγ and CEBPα while the level of the lipolytic gene (ATGL) is significantly (*P* < 0.05) downregulated (Fig 6C). Furthermore, as shown in Fig 6D, the results of the protein levels of the related pathway genes from the previous KEGG enrichment analysis revealed that the protein levels of Nox4, one of the NADPH oxidases, which plays a key role in intracellular ROS production in mammalian cells, and RAGE increased significantly (*P* < 0.05) while the level of p53 decreased significantly (*P* < 0.05). Based on this result, our subsequent research focused on the AGE-RAGE signaling pathway.

Based on the previous studies, which indicated that the AGE-RAGE signaling pathway may share a key role with ROS during the mediation of adipocyte differentiation, the antioxidant NAC was applied to inhibit ROS production in the primary adipocytes at the concentration of 10 mM for the first 2 days (days 0-2) of the standard differentiation protocol. As shown in Fig 7A, 2 days of NAC treatment cause significantly (*P* < 0.01) attenuated lipid accumulation to the normal level (compared with the control group) under the stimulus of TiO_2_ NPs, as detected using Oil Red O staining. Additionally, after adding NAC to the culture medium under the TiO_2_ NPs, the ROS measurement results revealed that NAC inhibited the level of ROS in the primary adipocytes with the fluorescence intensity of the DCF probe significantly reduced (Fig 7B). In addition, the protein levels of Nox4 and RAGE were significantly (*P* < 0.05) weakened after the addition of NAC to the treatment with TiO_2_ NPs (Fig 7C). In summary, these results suggest that TiO_2_ NPs accelerated lipid accumulation in the adipocytes accompanied with the upregulation of intracellular ROS levels and the expression of RAGE, but an application of the antioxidant NAC, in turn, attenuated lipid accumulation as well as the protein level of RAGE.

**Fig 7.**
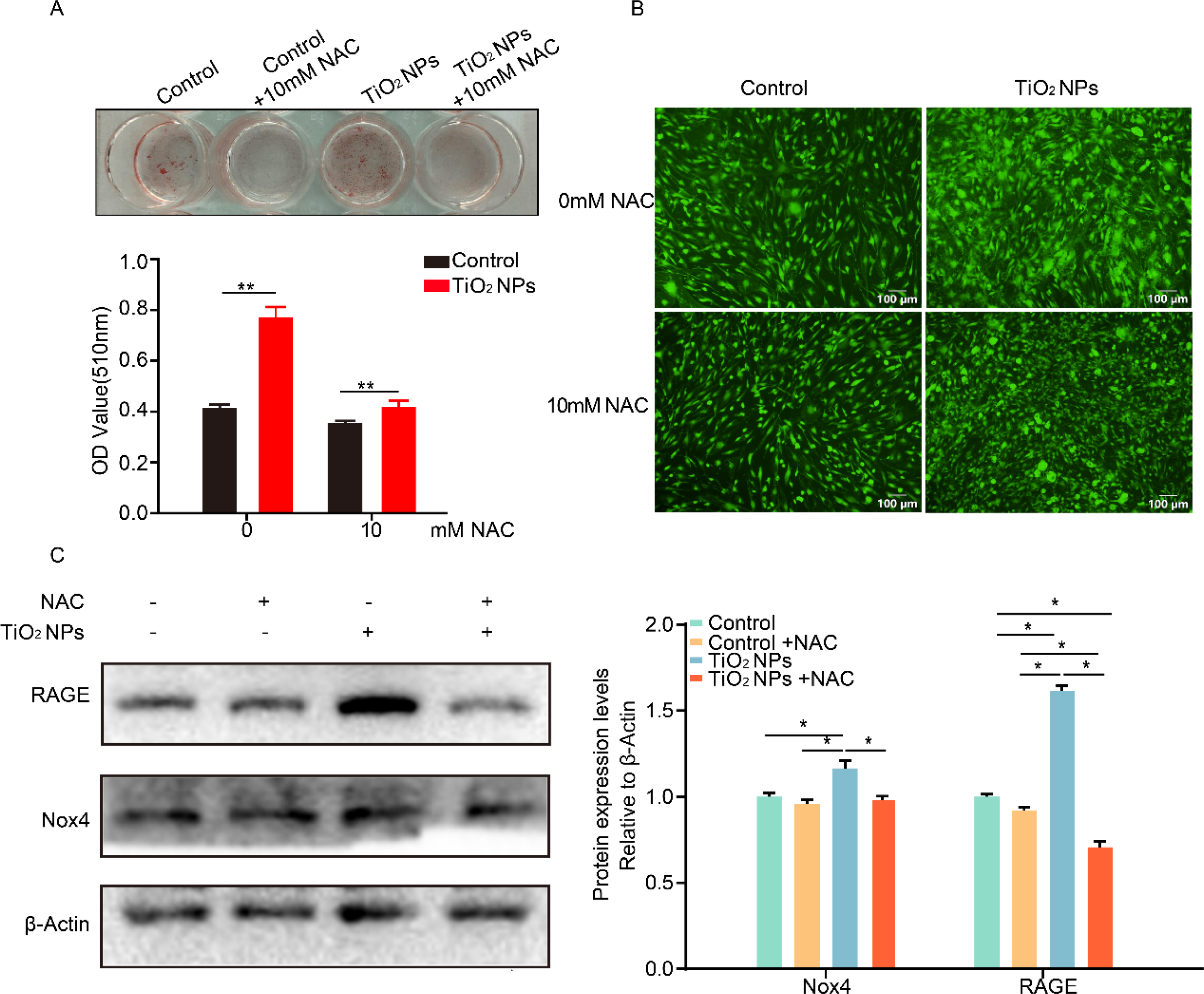
Effects of the antioxidant NAC on lipid accumulation induced by TiO_2_ NPs in the mice primary adipocytes. (A) Oil Red O staining diagram under different treatments. (B) Fluorescence images of intracellular ROS levels under different treatments. (C) Western blot of the related pathway genes in different treatments. Data represent mean ± standard error, (n = 3), \**P*<0.05, ** *P*< 0.01 and \*\*\**P* < 0.001 compared to the control group.

## 4. Discussion

With the development of science and technology, nanomaterials are widely used in various fields, and people cannot avoid contacting nanomaterials through various means. In the meantime, we particularly focused on the effect of TiO_2_ NPs from the digestive tract on the body. Our results indicated that TiO_2_ NPs exposure, at a dose of 100 mg/kg BW, will cause a blood glucose metabolism disorder, and an increase in blood glucose has nothing to do with weight and diet, which is consistent with the previous observations by Hu et al (Hu et al., 2016; Hu et al., 2018). More alarmingly, the significant increase in epididymal fat mass under the exposure to TiO_2_ NPs (Figs 2F and G), and the overexpression of the lipogenic genes (PPARγ, FABP4, and FASN) may contribute to abnormal fat deposition in the eWATs (Fig 2K), which suggests metabolic abnormalities in the body (Brestoff & Artis, 2015). Taken together, these data strongly suggested that TiO_2_ NPs cause damage to the glycolipid metabolism of the organism. Meanwhile, we chose 3T3-L1 preadipocytes as an *in vitro* model to explore the mechanism of TiO_2_ NPs during proliferation and differentiation of preadipocytes to mature adipocytes from the cellular level. Our results indicated that TiO_2_ NPs exposure to the adipocytes at a high concentration of (100 µg/mL) was sufficient to cause the induction of cell proliferation (Fig 3) and the enhancement of cell differentiation (Fig 4).

In addition, we used proteomic analysis to explore the mechanisms of TiO_2_ NPs to promote the differentiation of 3T3-L1 cells and identify the differentially expressed proteins. Many studies have shown that the accumulation of TiO_2_ NPs activates intracellular oxidative stress, with a massive release of ROS and lipid peroxides (Kong et al., 2022; Wu & Xie, 2016; Liu et al., 2010). Abnormal induction of ROS leads to dysregulation of other metabolic activities in cells, such as lipid metabolism and apoptosis. On this basis, we found that the AGE-RAGE signaling pathway may contribute to differentiation and lipid accumulation in the adipocytes, as well as ROS formation (Bierhaus & Nawroth, 2009; Monden et al., 2013; Unoki et al., 2007). The AGE-RAGE signaling pathway affects the level of inflammation and the function of adipose tissues by activating the activity of NADPH oxidase in cells (Song et al., 2014; Chen, Abell, Moon, & Kim, 2012; Coughlan et al.2009). Previous studies have shown that AGEs can directly promote the formation of free radicals, enhance the level and activity of NADPH oxidase, and weaken the cellular antioxidant system composed of glutathione peroxidase and Cu/ZnSOD (Paget, Lecomte, Ruggiero, & Wiernsperger, 1998). Studies on mouse adipose tissue showed that RAGE-mediated oxidative stress can lead to increased levels of the plasminogen activator inhibitor 1 and monocyte chemoattractant protein 1, both of which are related to the occurrence of obesity-related glucose intolerance (Unno et al., 2005; Uchida et al., 2004). *In vitro,* cultured adipocyte experiments showed that AGEs can induce NADPH oxidase to produce oxidative stress through RAGE, and induce the downregulation of leptin and adiponectin levels. Adiponectin itself can also affect its function through glycosylation, leading to the occurrence of insulin resistance (Frizzell et al., 2009). In addition, studies on 3T3-L1 preadipocytes have shown that the AGE-RAGE reaction can cause cellular insulin resistance by promoting the production of intracellular ROS (Unoki et al., 2007). Previous studies have shown that RAGE can mediate the production of intracellular ROS through Nox4, promote the generation of AGEs, and form the AGE-RAGE cycle (Malik & Kumar Mukherjee, 2022; Chen et al., 2012; Unoki et al., 2007; Hofmann et al.1999). Consistent with the results of previous studies, our findings revealed that the intake of TiO_2_ NPs will break the oxidation balance, caused an increased in ROS and promoted the production of AGEs, activated the receptor protein RAGE on the cell membrane, increased its expression, and then caused an increase in ROS. In addition, lipid overaccumulation in the primary adipocytes under the exposure to TiO_2_ NPs was impaired to the normal level with the application of the antioxidant NAC at a concentration of 10 mM (Lee et al., 2009), which was accompanied with the reduction in the level of ROS. Meanwhile, the level of RAGE in the adipocytes under the treatment of TiO_2_ NPs was significantly downregulated with the management of NAC.

In summary, exposure to TiO_2_ NPs could accelerate lipid overaccumulation and increase the level of ROS with the level of RAGE upregulated in the adipocytes. Notably, the application of the antioxidant NAC rescued the adipocytes from lipid overaccumulation by downregulating the level of RAGE, which indicates the potential therapeutic effect of exposure to TiO_2_ NPs on abnormal fat deposition.

## 5. Conclusions

In the present study, we revealed that exposure to TiO_2_ NPs (100 mg/Kg BW, 20 nm) accelerated abnormal fat deposition in the eWAT and disturbed the level of blood glucose subsequently in normal-fat diet mice. *In vitro* studies in the 3T3-L1 preadipocytes indicate that a high concentration of TiO_2_ NPs (100 µg/mL) induced cell proliferation and differentiation. Mechanistically, exposure of TiO_2_ NPs mediates ROS accelerated fat deposition through AGE-RAGE signaling pathway. However, the antioxidant NAC could impair lipid overaccumulation in the adipocytes to normal by downregulating the levels of RAGE in the adipocytes. Taken together, our research highlights the importance of reevaluating and comparing the effects of TiO_2_ NPs, especially those used in foods, on fat deposition, as well as providing a potential therapy for the complications of the disruption in lipid metabolism by TiO_2_ NPs.

## Author contributions

**Jingyi Liu** and **Liang Zhong** completed the thesis writing and main experiments. **Tiantian Yuan**, **Jiahao Chen** and **Yaxin Wang** helped feed mice and isolation of the primary adipocytes. **Jingchun Sun** provided a guide in cell culture. **Jingyi Liu**, **Liang Zhong**, and **Taiyong Yu** contributed to the experiment design. **Gongshe Yang** provided experimental platforms. **Gongshe Yang** and **Taiyong Yu** provided funding sources and overall ideas.

## Acknowledgments

This work was supported by the National Key R&D Program of China (2021YFD1301200).

---

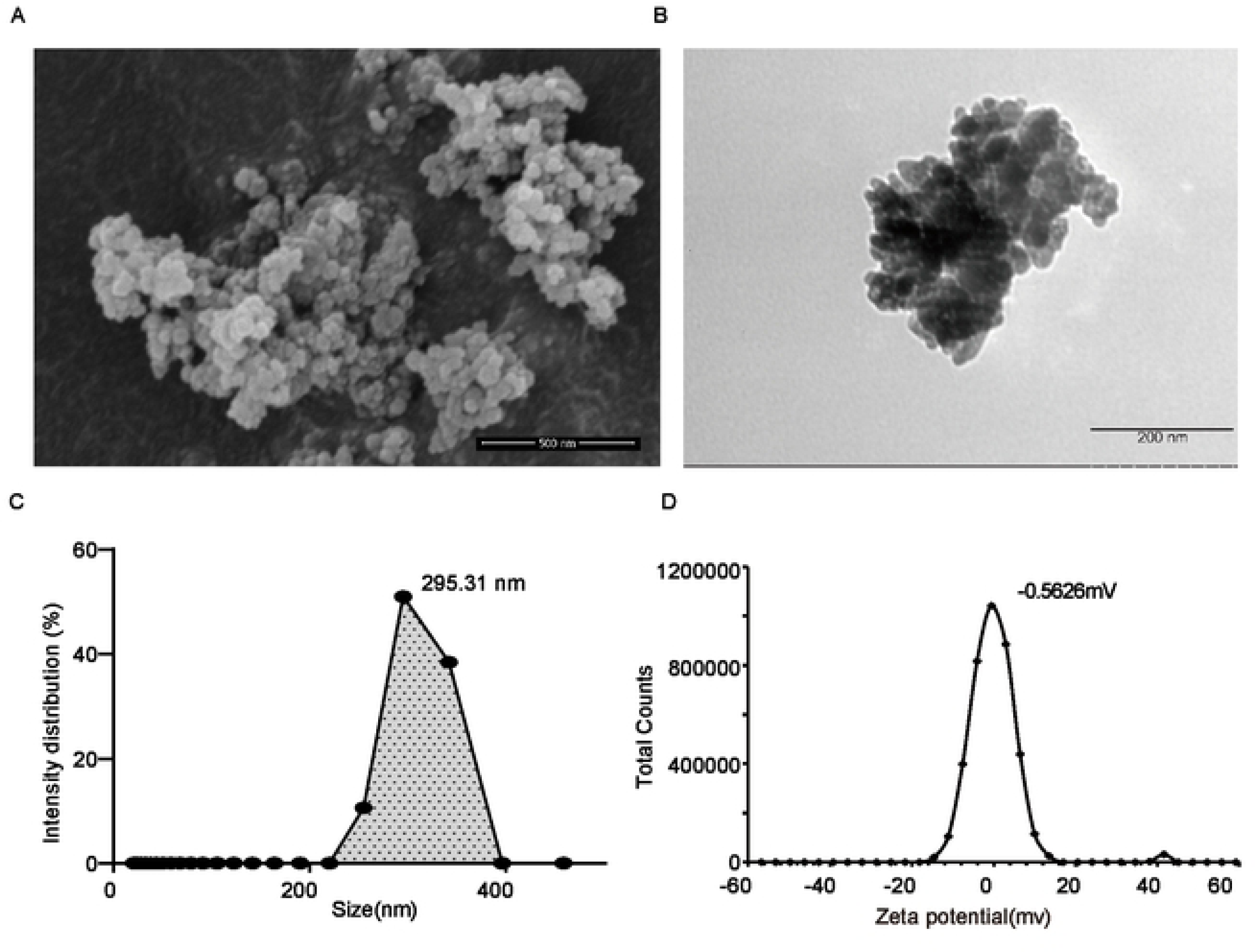

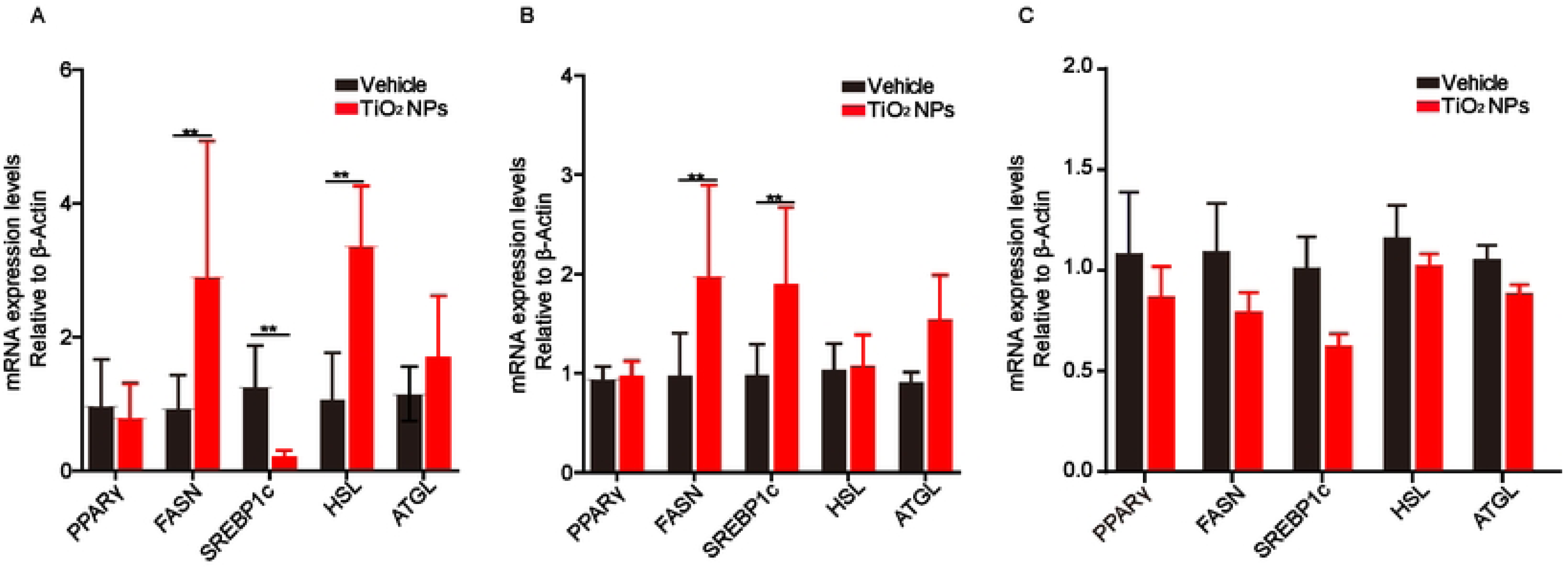

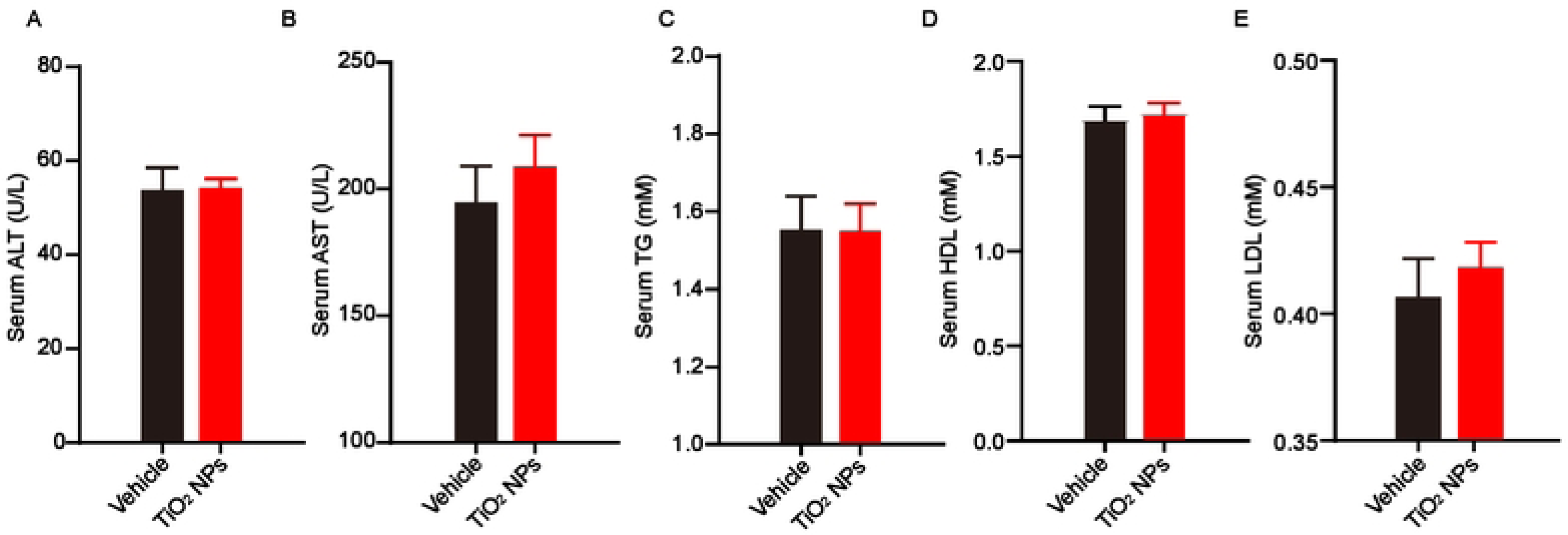

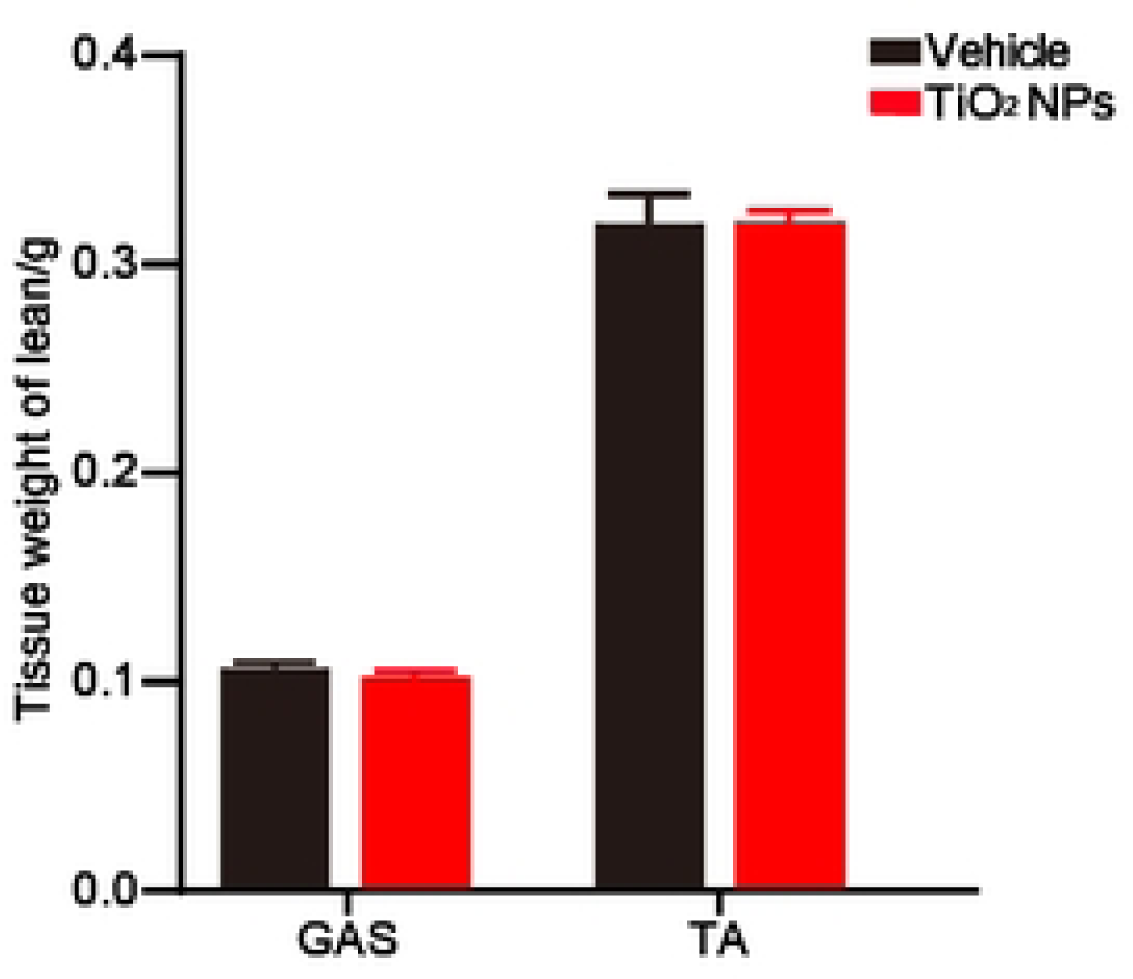

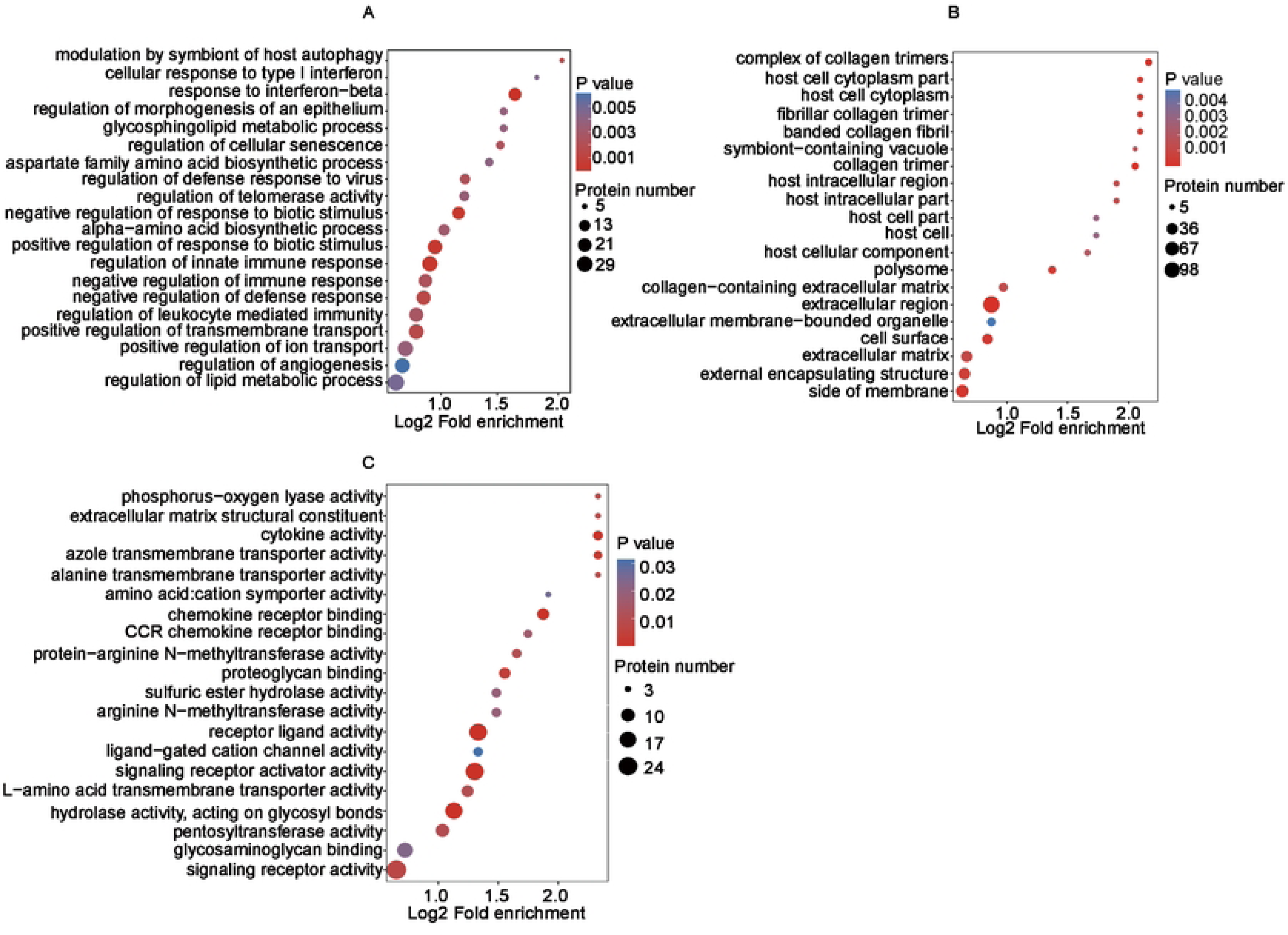

